# A Scalable Cross-Modal Workflow for Linking *In Vivo* Neural Activity to Neuronal Cell Types via Spatial Gene Profiling

**DOI:** 10.64898/2026.08.26.747400

**Authors:** Risa Yamazaki, Mark Eddison, Anja Payne, Greg Fleishman, Yuhan Wang

## Abstract

A central challenge in systems neuroscience research is to elucidate how distinct neuronal cell types coordinate to produce complex behaviors: an endeavor that requires linking their molecular identity, connectivity and activity patterns in behaving animals. Recent advances in single-cell RNA sequencing and spatial transcriptomics have revealed remarkable molecular diversity among neurons. However, most functional recording techniques, including *in vivo* two-photon calcium imaging, do not reveal the molecular cell identities of recorded cells. Although manual one-to-one matching between functional imaging and *post hoc* molecular profiling has been achieved for dozens to a few hundred neurons, these approaches are typically labor-intensive, difficult to scale, and capture only a limited fraction of cell types. Integrating these two modalities at large scale remains challenging, limiting our ability to fully understand the general logic of neural computation. Here we present a cross-modal workflow that integrates *in vivo* two-photon calcium imaging in behaving mice with Expansion-Assisted Iterative Fluorescence *In Situ* Hybridization (EASI-FISH), a thick-tissue spatial transcriptomic approach. As a proof of concept, we apply this workflow to the mouse dorsal hippocampus, a brain region with high neuronal density and small cell size where manual cell matching is impractical, making it a stringent test of the method’s accuracy and scalability. Using our approach, we achieved highly accurate alignment between *in vivo* neural activity and *ex vivo* molecular cell-type identity across hundreds to thousands of neurons, enabling direct mapping of neural dynamics to molecularly defined neuronal populations. We showed that longitudinal neural recordings followed by molecular mapping reveal distinct neural activity patterns that emerge as animals learn a spatial navigation task. By linking activity patterns to molecularly defined cell types, this method provides a framework for testing whether neural representations are organized through cell-type-specific coding or distributed population activity, offering insight into the computational logic by which neural circuits generate complex behaviors.

**MOTIVATION:** The mammalian brain contains an immense diversity of neurons exhibiting substantial morphological, functional, and molecular diversity. Understanding how this molecular heterogeneity supports neural computation and shapes behavior remains a central challenge in systems neuroscience. Yet, most approaches for recording neural activity provide limited access to molecular identity, making it difficult to directly relate functional dynamics to cell-type-specific features across many neurons in the same tissue. Here, we present a cross-modal workflow that recovers, aligns, and matches *in vivo* neural activity with targeted molecular profiling, enabling functional readouts and RNA profiling to be coupled at the individual cell level. This method provides a powerful platform for uncovering how molecularly defined neurons contribute to complex activity patterns and, ultimately, animal behavior.

## INTRODUCTION

The mammalian brain has a large diversity of cell types, comprising neurons that differ along molecular, morphological, anatomical, and functional axes ^1–8^. Understanding how this diversity supports neural computation and behavior remains a central question in neuroscience that has been constrained by the available genetic approaches. Recent advances in single-cell RNA sequencing and spatial transcriptomics have generated high-resolution catalogues of transcriptionally distinct neuronal populations across the mammalian brain ^9–17^. In parallel, *in vivo* two-photon calcium imaging ^18^, together with genetically encoded calcium indicators such as GCaMPs ^12,19,20^, enables longitudinal recording from large neuronal populations in behaving animals. However, these two areas have largely advanced separately: most functional-recording approaches do not resolve the molecular identity of recorded cells, limiting our ability to test whether neural representations are organized according to molecularly defined neuronal populations or distributed across molecularly diverse cell types.

Several approaches have begun to link *in vivo* activity with molecular identity. Cre- and Flp-dependent transgenic lines, often combined with viral tools, have been widely used to record from genetically defined neuronal populations ^21–23^. However, this strategy depends on the availability and specificity of driver lines and typically relies on a small number of marker genes, restricting analysis to a limited set of pre-selected cell classes. Other approaches that combine *in vivo* recording with *post hoc* cellular profiling provide complementary information but are often limited in throughput or by the number and diversity of neurons that can be profiled per experiment ^24,25^. More recently, multiplexed *in situ* hybridization-based spatial transcriptomic methods ^26–28^ have enabled gene-expression profiling at cellular resolution while preserving spatial context in intact tissue ^9–12,16,29–32^. Such methods provide a natural route for linking neural activity with molecular measurements at single-cell resolution. Building on this foundation, recent studies have combined *in vivo* functional imaging with *post hoc* spatial molecular profiling in the same tissue ^33–42^. Together, these advances have begun to reveal that molecular identity provides important context for interpreting functional heterogeneity across neurons, underscoring the need for broader adoption of these multimodal approaches across diverse neural systems.

Despite this progress, linking *in vivo* activity with *post hoc* molecular identity at scale remains technically challenging. First, tissue recovery can be difficult. Many workflows rely on thin tissue sections, which are vulnerable to damage and distortion during cryo-sectioning and downstream processing. In addition, small differences in sectioning angle or anatomical position can also compromise recovery of the *in vivo* imaged plane, resulting in partial or complete loss of the recorded population. These sources of variability reduce reproducibility across samples and limit the scalability of downstream registration and molecular profiling. Second, *in vivo* and *ex vivo* image volumes differ substantially in resolution, contrast, tissue texture, fluorescence patterns, and optical distortions. These modality-specific differences, together with tissue deformation during processing, make cross-modal registration challenging. This problem is further compounded in densely packed tissues, where individual cells are difficult to distinguish based on position and morphology alone. Consequently, manual one-to-one cell matching becomes impractical beyond a few hundred neurons, and small registration errors can lead to ambiguous or incorrect cell assignments.

An effective workflow for linking *in vivo* function with *post hoc* molecular identity should therefore meet several criteria. First, it should be reliable and straightforward to adopt. Because *in vivo* experiments require cranial surgery, behavioral training, and longitudinal recording, each sample represents a substantial experimental investment. Downstream tissue handling, imaging, and registration steps must therefore support reliable recovery of the recorded population while adding minimal complexity to standard *in vivo* imaging pipelines. Second, the workflow should be scalable. Growing evidence suggests that neural computation often depends on population-level activity patterns ^43–45^ that cannot be fully inferred from small numbers of neurons. Thus, molecular profiling should be able to scale with modern *in vivo* imaging approaches, including large field-of-view and multi-plane recordings that capture hundreds to thousands of neurons.

To address these challenges, we developed a scalable cross-modal workflow that integrates large field-of-view two-photon mesoscope ^46^ imaging during behavior with thick-tissue Expansion-Assisted Iterative Fluorescence *In Situ* Hybridization, or EASI-FISH ^32^ for *post hoc* spatial gene-expression profiling in the same neurons. This workflow is designed to integrate with standard *in vivo* calcium imaging pipelines by using the native GCaMP fluorescence signal acquired during routine recording as the cross-modal anchor and avoid the need for additional fiducial labeling. For *post hoc* molecular profiling, we use thick tissue sections (300-500 µm) prepared with a standard vibratome. These sections preserve the *in vivo* imaged volume within a single slice, reducing sensitivity to sectioning angle and minimizing the need to reconstruct volumes from serial sections. In addition, EASI-FISH is compatible with thick tissue volumes and large field-of-views, enabling molecular profiling of large numbers of neurons recovered from *in vivo* imaging sessions. To make cell matching practical at this scale, we developed a semi-automated cross-modal alignment strategy that reduces the required manual effort. By moving beyond manual one-to-one matching, which becomes impractical at large population scales, this approach supports matching across hundreds to thousands of neurons per animal. As proof of principle, we applied this workflow to dorsal CA1 pyramidal neurons, a stringent test case where neurons are small, densely packed, and difficult to match by hand. When combined with longitudinal calcium imaging, this workflow recovered close to one thousand matched neurons from a single animal and enabled cell-resolved analysis of how molecular features relate to *in vivo* activity during behavior. Together, this workflow provides a practical approach for relating molecular identity to neural activity in large populations of neurons recorded during behavior. More broadly, the same principles may be adapted to other tissues and physiological contexts in which functional imaging can be paired with *post hoc* molecular profiling, providing a framework for linking gene expression with cellular function beyond the brain.

## RESULTS

### Overview of the integrated molecular and functional profiling workflow

We first implemented this workflow by combining longitudinal *in vivo* calcium imaging ^47^ with *post hoc* thick-tissue spatial gene-expression profiling in the same brain volume (**Figure 1A**). Neuronal activity was recorded and tracked during behavior using large field-of-view two-photon mesoscope imaging ^46^. At the end of the imaging sessions, and prior to transcardial perfusion, an *in vivo* z-stack of the native GCaMP fluorescence signal was acquired on the same two-photon microscope. The recovered brain was then horizontally sectioned and processed with EASI-FISH ^32^ for image-based spatial gene-expression profiling (**Figure 1B**). Unlike approaches that require additional staining or imaging steps to generate fiducial or consensus landmarks for *in vivo*-to-*ex vivo* alignment, this approach repurposes the native GCaMP fluorescence signal already acquired during standard *in vivo* calcium imaging experiments. Thus, no additional fiducial labeling or specialized tissue treatment is required. This design enables retrospective recovery of tissue samples for spatial gene-expression profiling while adding minimal complexity to existing *in vivo* and *ex vivo* workflows.

**Figure 1.**
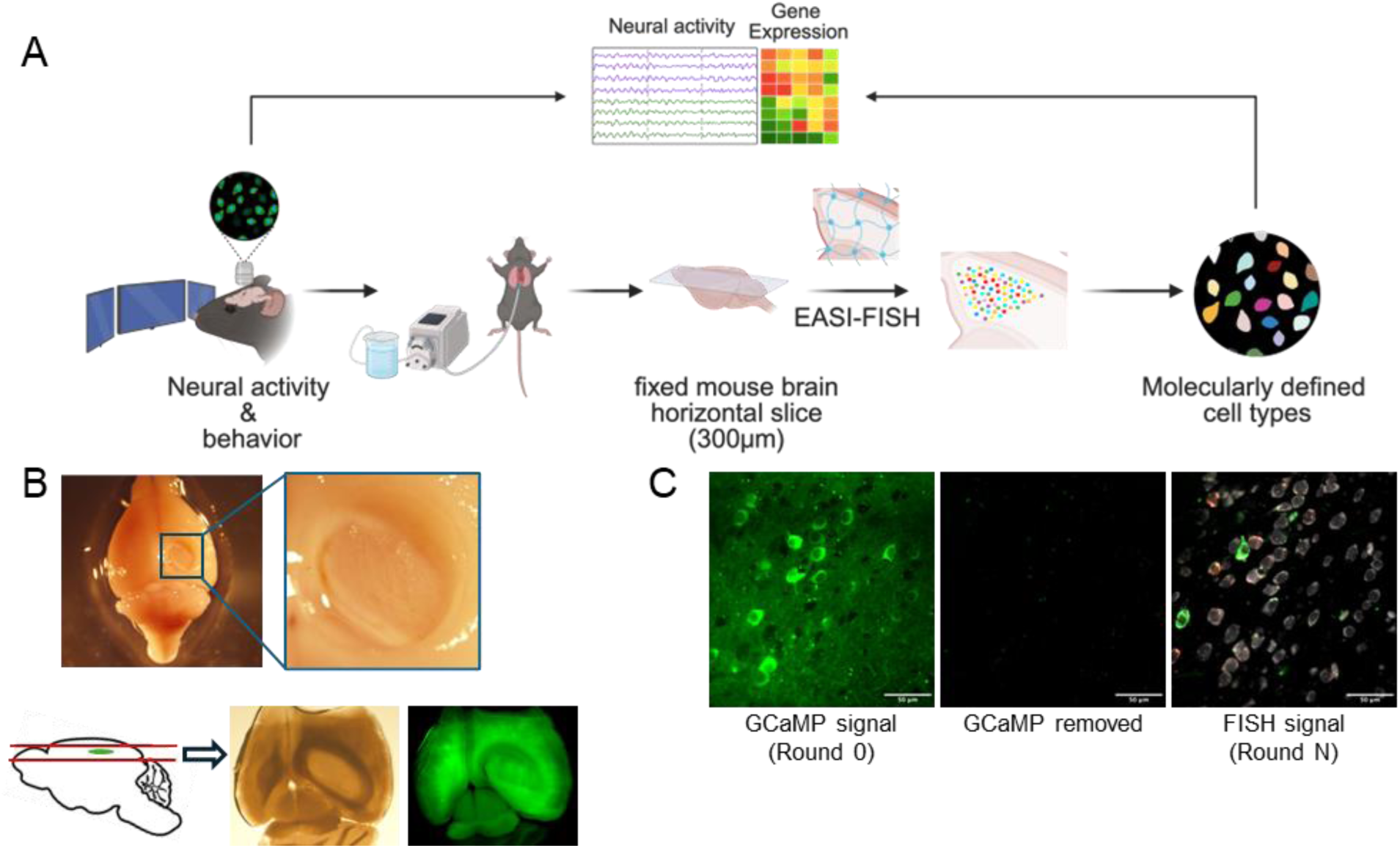
Integration of *in vivo* two-photon calcium imaging with *ex vivo* spatial gene expression profiling. (**A**) Overview of the experimental workflow. Mice undergo head-fixed behavioral training with *in vivo* two-photon calcium imaging. After transcardial perfusion, brain tissue is recovered for spatial gene expression profiling using Expansion-Assisted Iterative Fluorescence *In Situ* Hybridization (EASI-FISH). This approach allows neural activity to be measured in molecularly defined neuronal populations. (**B**) Tissue recovery workflow for registration and *ex vivo* spatial gene expression profiling. Top left: Representative image of a successfully perfused and recovered mouse brain. The boxed region marks the region of interest (ROI) beneath the cranial window. Top right: zoomed-in view of the ROI containing the tissue beneath the cranial window. Bottom left: side-view schematic showing the horizontal sectioning strategy used to recover the ROI. The brain was oriented to align the sectioning plane to the *in vivo* imaging plane. The green oval denotes the ROI. Bottom middle and right: Representative tissue section containing the recovered ROI. (**C**) *Ex vivo* GCaMP imaging for registration and subsequent signal removal for EASI-FISH. Left: representative GCaMP image from Round 0. Middle: complete removal of GCaMP signal after round 0. Right: FISH signals (shown in green and red) and cytoDAPI (shown in grayscale) from subsequent imaging rounds after GCaMP removal. Scale bars: 50 µm.

After transcardial perfusion, thick tissue sections (typically 300 µm), were collected for EASI-FISH. Because EASI-FISH is compatible with thick sections, this approach increases tolerance to differences in sectioning angles between the *in vivo* and *ex vivo* preparations. Compared with thin-section methods, which often require extensive cell matching across serial sections and volumetric reconstruction, the thick-section strategy minimized the loss of *in vivo* imaged neurons, maximized their recovery for gene expression profiling, and streamlined downstream *ex vivo* processing and alignment. We found that modified EASI-FISH protocol (see STAR method) effectively preserved GCaMP fluorescence. During *ex vivo* GCaMP imaging, we also acquired the cytoDAPI signal generated after DNase treatment, as previously described. After acquiring a GCaMP image stack for *in vivo*-to-*ex vivo* registration (Round 0, **Figure 1C, left**), we used a detergent (SDS) and proteinase K-based protocol to eliminate GCaMP fluorescence, thereby freeing the GCaMP channel for FISH imaging (**Figure 1C, middle**). This approach reliably eliminated GCaMP fluorescence from both transgenic and viral sources while preserving RNA integrity for downstream molecular analysis (**Figure 1C, right**). The *ex vivo* preserved GCaMP stack serves as a bridging imaging round for registration to the *in vivo* calcium imaging volume, whereas the cytoDAPI channel provided a reference for alignment with subsequent FISH imaging rounds (**Figure 2A**). The cytoDAPI signal was also used for cell body segmentation and transcript assignment. The resulting *ex vivo* segmentation masks enabled two complementary approaches for linking molecular and functional datasets: they could either be matched to *in vivo* ROIs identified by standard calcium imaging analysis pipelines or transformed into the *in vivo* coordinate space and used directly to extract neural activity traces from the *in vivo* imaging data.

**Figure 2.**
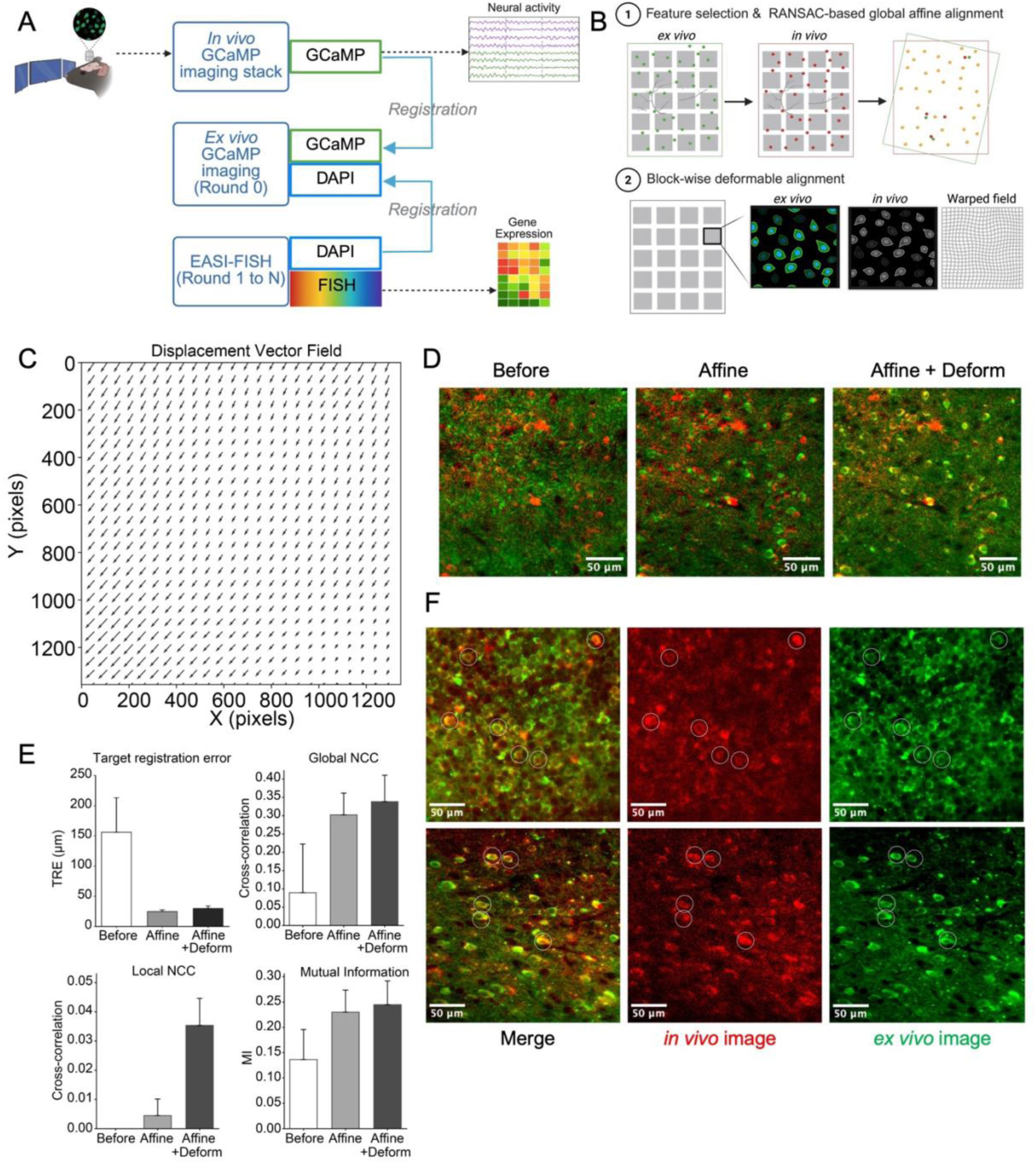
Registration procedures. (**A**) Strategy for *ex vivo*-to-*in vivo* registration. *Ex vivo* GCaMP imaging in Round 0, together with cytoDAPI, serves as a bridging imaging round that links *in vivo* neural activity data to gene expression measurements acquired in subsequent EASI-FISH rounds (rounds 1 to N). (**B**) Schematic overview of the two-step registration workflow. (**C**) Representative transformation used to warp the *ex vivo* image into the *in vivo* imaging coordinate space. (**D**) Representative overlays of *in vivo* (red) and *ex vivo* (green) images before registration (Before), after the first step of alignment (Affine), and after the two-step alignment (Affine + Deform). Scale bars: 50 µm. (**E**) Registration performance quantified using target registration error (TRE), global and local normalized cross-correlation (NCC), and mutual information (MI). Data are shown as mean ± standard deviation. (**F**) Representative images showing corresponding cells identified in the *in vivo* GCaMP image (red) and the *ex vivo* GCaMP image (green), after the two-step registration workflow. Circles highlight examples of matched cells across the two images. Scale bars: 50 µm.

### Two-step image registration to align *in vivo* and *ex vivo* image volumes

Registration of *in vivo* and *ex vivo* imaging modalities is challenging because the two datasets differ substantially in resolution, contrast, tissue texture, fluorescence signal patterns, and tissue geometry. *In vivo* two-photon images are acquired with relatively low axial resolution and contain activity-dependent GCaMP signal. In contrast, *ex vivo* EASI-FISH volumes are acquired at higher, near-isotropic resolution and contain a broader GCaMP fluorescence pattern. These differences are further compounded by modality-specific optical and physical distortions, including optical aberrations, field curvature, tissue scattering and tissue deformation during processing, which can alter the apparent geometry and scaling of neuronal structures. Although manual approaches have been used, they are labor intensive, error-prone, and typically recover only small populations of neurons, mainly in regions with sparse cell distributions. In densely packed brain regions, high-precision manual alignment becomes particularly difficult because individual cells are challenging to distinguish based on position and morphology alone, and small registration errors can lead to ambiguous or incorrect cell assignments.

To overcome these challenges, we developed a two-stage registration pipeline to align the *ex vivo* image volume, used as the moving image, to the *in vivo* image volume, used as the fixed image (**Figure 2B**). In the first stage, we estimated a global affine transformation using manually annotated corresponding landmarks, typically 30-50 pairs, based on anatomical and tissue features (e.g., blood vessels). Landmark pairs were defined in physical coordinates, and the affine transformation matrix was estimated using random sample consensus (RANSAC) to reject outliers and compute an optimal 3D affine mapping. This global registration step corrected global differences in translation, rotation, orientation, and scaling between the two image volumes. In the second stage, we performed B-spline based block-wise deformable registration to correct local, nonlinear tissue distortions. The deformable registration was optimized by gradient descent using ANTs neighborhood correlation (ANC) as the similarity metric. To improve computational efficiency, the image volume was divided into spatial blocks that could be registered in parallel. This distributed strategy enabled scalable registration of large volumetric datasets that would otherwise be difficult to process as a single volume.

As a stringent test case, we applied this workflow to dorsal CA1 of the mouse hippocampus, where pyramidal neurons are densely packed and manual one-to-one cell matching is impractical. The two-stage registration approach performed consistently across samples and corrected both global and local mismatches. The estimated deformation fields were physically plausible and did not introduce obvious topological artifacts (**Figure 2C**). Cellular-level correspondence improved progressively after each registration stage. The global affine transformation brought the two image volumes into approximate alignment, whereas the local deformable registration further refined cell-level correspondence. Before registration, individual neurons visible in both modalities were spatially misaligned (**Figure 2D**, left: Before). The global affine transformation brought the two images into approximate alignment, but residual local mismatches remained (**Figure 2D**, middle: Affine). The local deformable registration further refined cell-level alignment, where corresponding cell bodies in the two images were spatially overlaid (**Figure 2D**, right: Affine + deform).

We quantified registration performance across hippocampal samples using complementary metrics, including target registration error (TRE, on holdout landmarks), as well as intensity-based global and local normalized cross-correlation (NCC), and mutual information (MI) (**Figure 2E**). The affine registration step substantially reduced TRE and improved global NCC and MI, indicating improved overall alignment between the two volumes. The deformable registration step further improved local NCC, consistent with correction of residual local nonlinear distortions. These results indicate that affine registration provides robust global alignment, while deformable registration is required for local cell-level refinement. Importantly, the pipeline improved registration in both regions with moderate cell density and regions with very high cell density (**Figure 2F** and **Video S1**), indicating that the approach is robust across brain regions.

After establishing *in vivo*-to-*ex vivo* registration, we applied the resulting transformation to warp the *ex vivo* segmentation mask into the *in vivo* coordinate space. To generate the *ex vivo* segmentation mask, we used Starfinity ^32^, a deep learning cell segmentation method based on *Stardist* ^48,49^. We found that the model we previously trained on other brain regions generalized well to hippocampal neurons and produced accurate cell body segmentations in dorsal CA1, despite differences in cell size, density, and morphology (**Figure 3A**). We next matched *ex vivo* EASI-FISH cell masks to *in vivo* calcium imaging ROIs. After motion correction and ROI extraction using Suite2p ^50,51^, we identified candidate matches by detecting overlapping ROI pairs between the transformed *ex vivo* mask and the *in vivo* Suite2p mask at the corresponding z-plane, to ensure compatibility with a widely used calcium imaging analysis pipeline. One-to-one ROI assignments were determined from two complementary criteria, shape similarity (measured by centered intersection-over-union or cIoU), and spatial proximity (measured by Euclidean centroid distance) (**Figure 3B-C**, see STAR Methods). A logistic regression on these two features was fit to manually labeled candidate matches, and the predicted probability threshold was selected based on maximum Youden’s index. Using this approach, we recovered 77% of *in vivo* imaged neurons (Precision: 0.891, 95% CI: 0.804-0.978), with the highest recovery rate in the center of the field-of-view. After cell-to-cell correspondences were established, neuronal activity traces and gene expression profiles were extracted from the same individual neurons (**Figure 3D** and **Video S2**).

**Figure 3.**
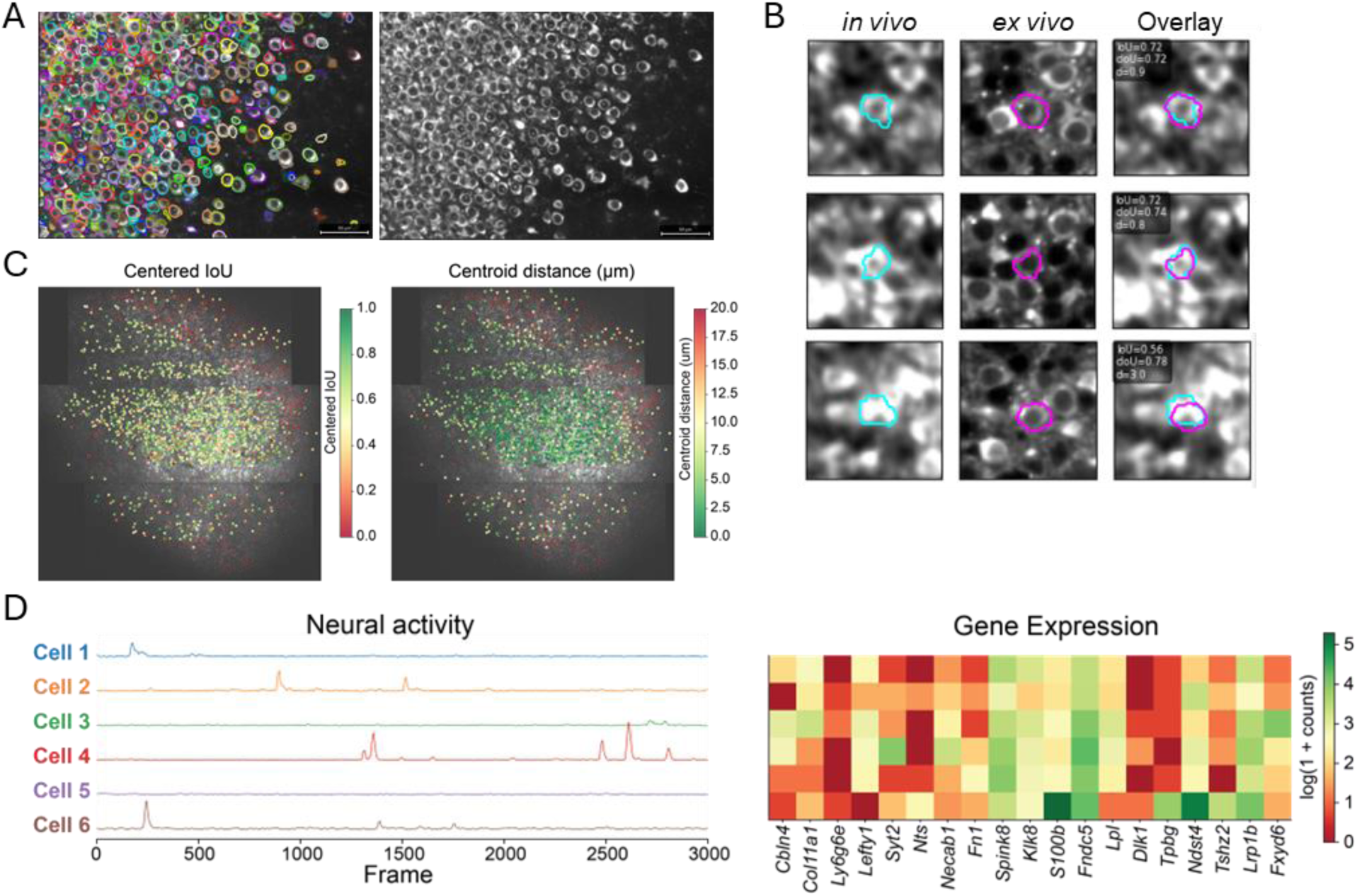
Cell-cell matching between *in vivo* and *ex vivo* datasets. (**A**) Representative image showing *Starfinity*-based cell segmentation on *ex vivo* images from the dorsal hippocampus. Scale bars: 50 µm. (**B**) Examples of three matched neurons based on *in vivo* and *ex vivo* cell masks. Left: *in vivo* GCaMP image with an example cell mask. Middle: Corresponding *ex vivo* cell mask overlaid on the *ex vivo* image. Right: Overlay of the matched *in vivo* and *ex vivo* masks. (**C**) Quantification of cell-matching quality. Left: Centered intersection-over-union (cIoU) values for *in vivo* ROIs. Right: Centroid distances between *in vivo* ROIs and their matched *ex vivo* ROIs. Both shown overlaid on the *in vivo* GCaMP image. (**D**) Representative neural activity traces and gene expression profiles from the same neurons after *in vivo*-to-*ex vivo* cell-cell matching.

### Molecular profiles reveal distinct activity and spatial coding properties in dorsal CA1 neurons

We next leveraged the matched molecular and functional datasets to investigate the relationship between gene expression and neural activity in the dorsal hippocampus. Hippocampal pyramidal neurons exhibit well-established molecular ^15,17,52–56^, connectivity ^57–59^, and functional ^60–64^ heterogeneity along multiple anatomical axes, including the deep-superficial ^65–67^, proximal-distal ^60,68^, and dorsal-ventral ^69–72^ axes, yet these anatomical, molecular, and functional dimensions have largely been examined in separate experimental contexts. As a result, the extent to which molecular programs predict the activity patterns of individual dorsal CA1 neurons during behavior remains poorly understood. To address this, we performed longitudinal two-photon calcium imaging in animals trained on a previously described virtual-reality two-alternative cue-delay-choice task^47^ (**Figure S1A**), followed by EASI-FISH on 20 marker genes selected to capture known anatomical and molecular heterogeneity within the dorsal CA1 and subiculum (**Figure 4A**). Marker gene expression analysis identified six molecularly defined neuronal populations, including three clusters from the dorsal CA1 (C1, C2, C3) and three clusters from the dorsal subiculum (C4, C5, C6) (**Figure 4B**). We focused the analysis on the two major dorsal CA1 clusters (C1 and C2), which were well represented within the matched *in vivo*-to-*ex vivo* dataset. Comparison of the overall functional properties revealed different spatial coding properties between C1 and C2. C1 neurons encode higher spatial information and have greater spatial coding stabilities between sessions as well as between trials (**Figure 4C**).

**Figure 4.**
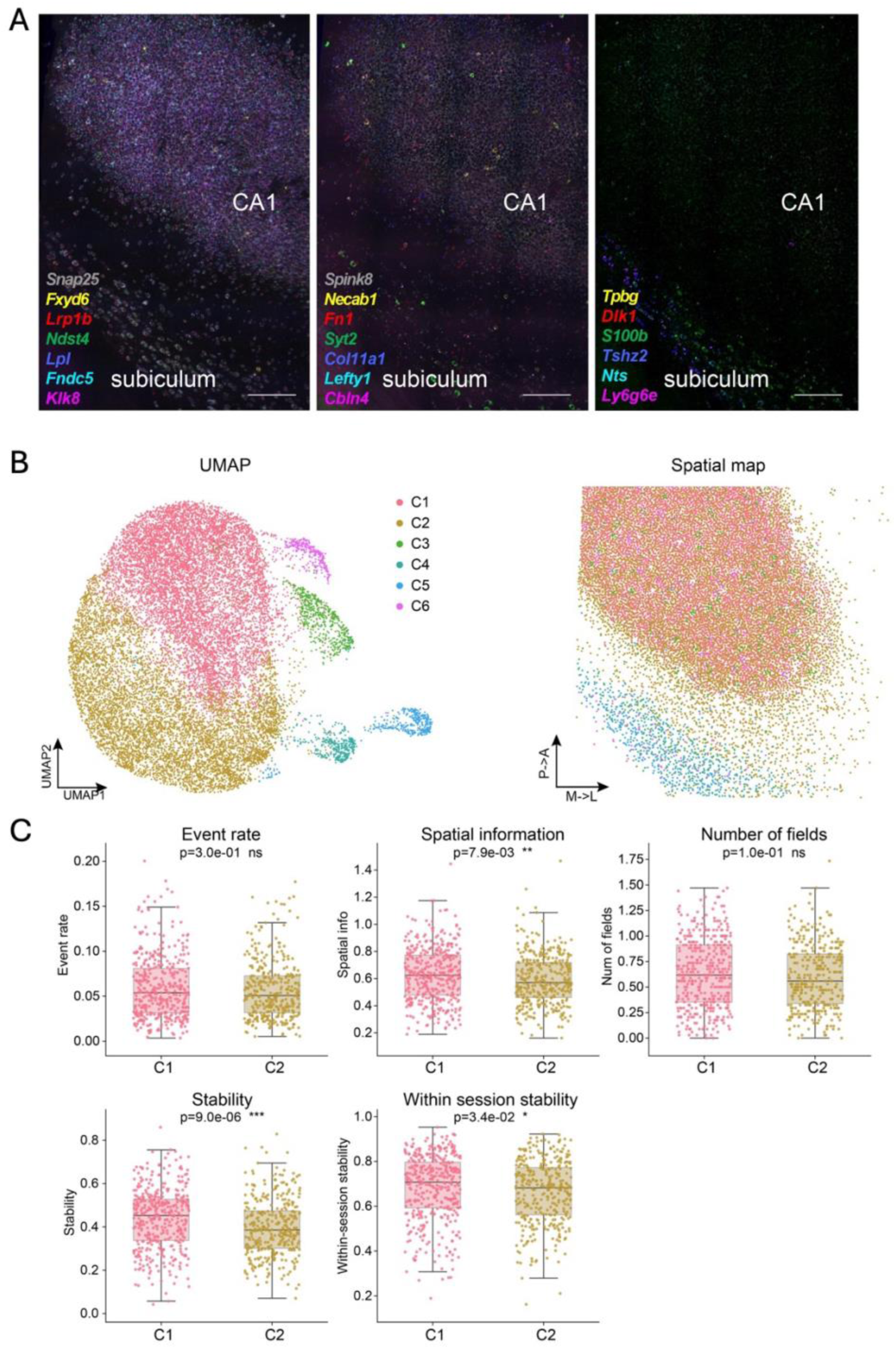
Molecular and functional characterization of EASI-FISH profiled hippocampal neurons. (**A**) Representative EASI-FISH images showing expression of 20 marker genes profiled in the mouse dorsal hippocampus. Scale bars: 200 µm. (**B**) UMAP (left) and spatial maps (right) of profiled hippocampal neurons, colored by molecular cluster. For the spatial map, anatomical orientation is indicated by arrows showing the posterior-to-anterior (P->A) and medial-to-lateral (M->L) axes. The length of each arrow on the spatial map corresponds to 200 µm. (**C**) Comparison of functional properties between molecular clusters C1 and C2, including event rate, spatial information, number of place fields, and spatial coding stability (across sessions and within sessions). Data are shown as mean ± standard deviation.

Because CA1 pyramidal neurons are thought to vary along continuous gene-expression gradients rather than form strictly discrete cell types ^53^, we performed principal component analysis (PCA) on the gene-expression matrix to capture continuous molecular variation across dorsal CA1 neurons. PC1 explained 16% of the total variance and captured a continuous molecular gradient that covaried with position along both the proximal–distal and superficial– deep axes of the CA1 pyramidal layer (**Figure 5A**). This gradient was associated with opposing expression of *Ndst4* and *Col11a1* at one end and *Klk8*, *Lefty1*, and *Lpl* at the other (**Figure 5B**). We next asked whether *in vivo* functional properties varied systematically along this molecular axis. Spatial coding properties changed across the gradient: neurons located more proximally and superficially, corresponding to lower *Ndst4* and *Col11a1* expression, carried more spatial information, exhibited higher within-session place map stability, and maintained more stable place fields across sessions (**Figure 5C**). Finally, we asked whether the association between the molecular gradient and spatial coding varied across learning. To test this, we calculated the Spearman rank correlation between PC1 and each spatial coding metric separately for each training stage: pre-training, early training, middle training, and late training (see STAR Methods). We then compared the resulting correlation coefficients across stages. We found that the magnitude of these molecular-functional correlations, as indicated by the correlation coefficient (ρ), varied across training, with the strongest correlation observed during the active learning phase (early and middle training sessions) (**Figure S1D**). These results suggest that molecularly heterogenous neurons are most functionally divergent during the period when animals are actively learning the spatial task. Together, these findings support the integration of EASI-FISH-based molecular profiling with longitudinal neural activity recordings and highlight the utility of this approach for uncovering biologically meaningful relationships between gene expression and neural activity patterns.

**Figure 5.**
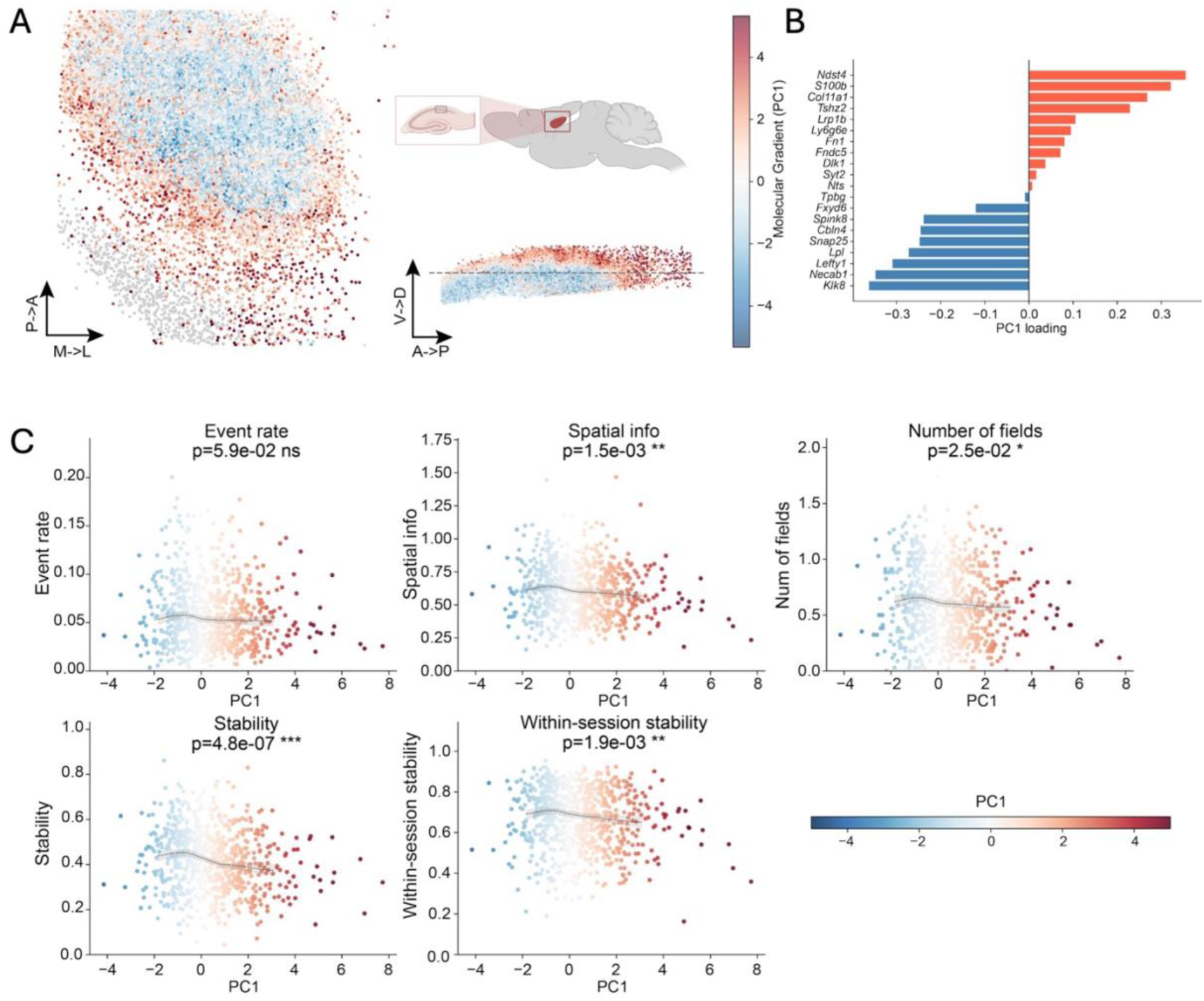
EASI-FISH analysis along a molecular gradient. (**A**) Spatial maps colored by the molecular gradient, defined by PC1. Left: Horizontal section view. Right: Sagittal view. For the spatial map, anatomical orientation is indicated by arrows. P->A: posterior-to-anterior, M->L: medial-to-lateral. V->D: ventral-to-dorsal. The length of each arrow corresponds to 200 µm. The dotted line in the sagittal view (right) indicates the *in vivo* imaging plane. (**B**) Loadings of marker genes on PC1. (**C**) Associations between functional properties and the molecular gradient, including event rate, spatial information, number of place fields, and spatial coding stability across sessions and within sessions. Each point represents a cell and is colored by its PC1 score. Spearman’s rank correlation was used to assess associations. Trend curves (LOESS smoothers with 95% confidence intervals) are shown for visualization only.

## DISCUSSION

In this work, we present a scalable experimental and computational workflow for integrating *in vivo* calcium imaging during head-fixed behavior with *post hoc* FISH-based molecular cell type mapping. This approach minimizes additional tissue handling and imaging steps by repurposing the existing GCaMP fluorescence signal for *in vivo*-to-*ex vivo* alignment, enabling retrospective molecular profiling of tissues collected after *in vivo* recordings without requiring additional fiducial labeling or specialized sample preparation. We further adapted EASI-FISH for intact thick tissue sections containing the *in vivo* imaged volume, thereby simplifying tissue recovery and avoiding extensive sub-sectioning and volumetric reconstruction. In addition, we developed a cross-modality registration strategy to align *in vivo* and *ex vivo* datasets, which differ substantially in resolution, contrast, optical distortions, and tissue geometry. This two-stage registration pipeline, consisting of coarse global affine alignment followed by local deformable refinement, reduces manual labor while preserving cellular-level alignment accuracy. Since the workflow relies on broadly available imaging signals and generalizable registration principles, it should be adaptable to other brain regions and experimental preparations with similar tissue recovery and imaging procedures. Together, these features substantially simplify the experimental and computational workflow and make the approach accessible for routine implementation.

As proof of principle, we applied this workflow to the dorsal hippocampus after large field-of-view two-photon mesoscope imaging. Mesoscope imaging enables simultaneous recording from thousands of neurons across broad cortical or hippocampal areas, providing the scale needed to relate molecular diversity to population-level neural activity. However, the large number of recorded neurons also makes manual cell matching impractical, particularly in dorsal CA1, where neurons are densely packed within the compact pyramidal cell layer. By applying our registration pipeline to this challenging preparation, we demonstrate that molecular and functional information can be linked at cellular resolution in a densely organized brain region. Combining EASI-FISH with longitudinal calcium imaging across learning revealed associations between molecular variation in CA1 pyramidal neurons and differences in spatial coding during behavior. Although this study sampled only a small portion of dorsal CA1, these results are broadly consistent with previous work showing that proximal and superficial CA1 neurons exhibit more stable spatial coding ^60,68,73^, potentially reflecting selective inputs to these regions ^59,63,65,74^. Together, this approach provides a framework for linking molecular variation to population coding, neural activity dynamics, and behavior.

Several extensions of this workflow should further increase its utility. In the present study, *in vivo* imaging was performed primarily in a single imaging plane. The workflow should readily support future extension to volumetric calcium imaging *in vivo*, which would enable larger-scale recovery of molecular and functional properties across three-dimensional neuronal populations. Additionally, the current implementation performs cell-to-cell matching between *ex vivo* EASI-FISH segmentation masks and *in vivo* ROIs generated by standard calcium imaging analysis pipelines, such as Suite2p. This design makes the workflow compatible with widely used *in vivo* analysis approaches. However, *ex vivo* segmentation masks may also be transformed into the *in vivo* coordinate space and used directly for neural activity extraction. Because *ex vivo* images are acquired at higher resolution and provide sharper cell-body boundaries, this strategy may improve cell recovery rates and reduce segmentation ambiguity in calcium data in future implementations. This workflow represents an important step toward linking gene expression, neuronal activity, and behavior in the same identified cells. Applying this framework across brain regions, behavioral paradigms, developmental stages, and disease models will reveal how molecular programs give rise to functional phenotypes and how these relationships are reshaped by experience or pathology.

## RESOURCE AVAILABILITY

### Lead contact

Further information and requests for resources and reagents should be directed to and will be fulfilled by the lead contact, Yuhan Wang

### Materials availability

This study did not generate new unique reagents.

### Data and code availability

- All original code is deposited at https://github.com/Y-Wang-Lab/2p_EASI-FISH-paper and is publicly available at the date of publication.
- Any additional information in this paper will be shared by the lead contact upon request.

## ACKNOWLEDGMENTS

We thank Cornell University College of Human Ecology and Division of Nutritional Sciences as well as Howard Hughes Medical Institute Janelia Research Campus for funding this work. We thank Dr. Nelson Spruston and his lab for support, resources and advice that enabled this work. We thank Dr. Scott Sternson, Dr. Paul Tillberg, and their labs for discussions on the project; Michalis Michaelos, Tyra Hay, and Rachel Gattoni for behavior training; Monique Copeland, Ben Foster and Amy Hu for histology support; Janelia Scientific Computing and High-Performance Computing team, Vivarium staff at Janelia Research Campus and Cornell University for animal support.

## AUTHOR CONTRIBUTIONS

Conceptualization and supervision: Y.W.; Methodology: Y.W., R.Y., M.E., A.P., G.F.; Investigation: Y.W., R.Y., M.E., A.P., G.F.; Writing: Y.W. and R.Y. with input from all co-authors.

## SUPPLEMENTAL INFORMATION

**Supplemental Figure 1.**
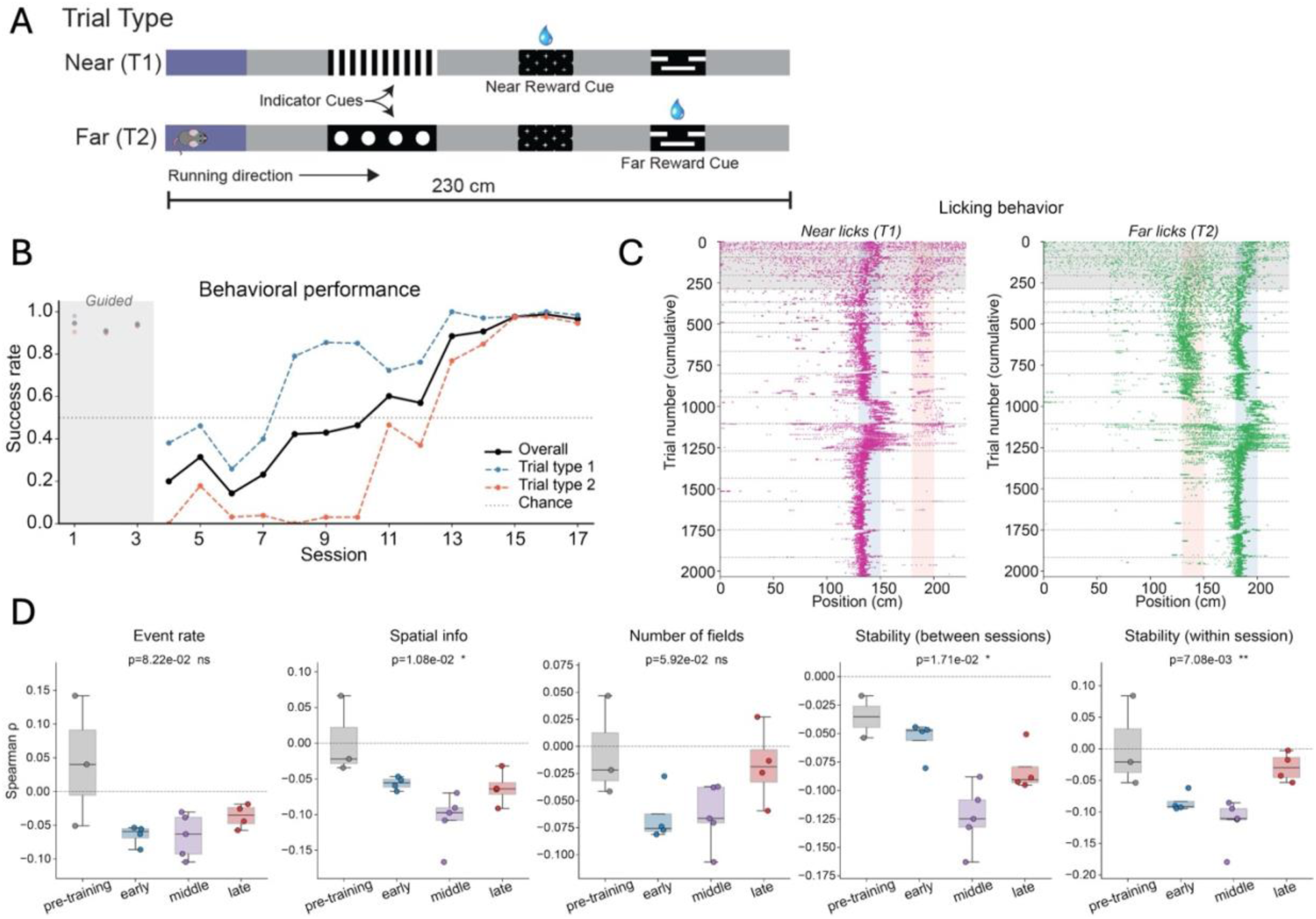
Molecular gradients are associated with neuronal properties during learning. (**A**) Schematic illustration of the behavioral task: Trial type 1 (T1, Near), with the reward zone located near the indicator cue and trial type 2 (T2, Far), with the reward zone located farther along the track. (**B**) Representative behavioral performance from one animal. (**C**) Licking behavior along the track across learning trials. The x-axis indicates track position, and each row represents a trial. Left: Trial type 1 (T1) and right: Trial type 2 (T2). (**D**) Strength of the molecular-functional association across learning, quantified as the Spearman rank correlation (ρ) between PC1 and each functional metric (event rate, spatial information, number of place fields, and within- and across-session spatial coding stability) at four training stages. Data are shown as mean ± standard deviation.

**Video S1.** Representative video showing the cross-modal alignment of *in vivo* (red) and *ex vivo* (green) GCaMP signals.

**Video S2.** Representative video showing (Top) EASI-FISH images registered to *in vivo* calcium imaging data to visualize neural activity and gene expression in the same neurons, and (Bottom) the corresponding *in vivo* (red) and *ex vivo* (green) GCaMP signals.

## STAR★METHODS

### KEY RESOURCES TABLE

| REAGENT or RESOURCE | SOURCE | IDENTIFIER |
| --- | --- | --- |
| **Chemicals, peptides, and recombinant proteins** |  |  |
| 16% Paraformaldehyde | Electron Microscopy Sciences | Cat. # 15710 |
| DPBS (calcium, magnesium) | Thermo Fisher Scientific | Cat. # 14040117 |
| 10× PBS | Thermo Fisher Scientific | Cat. # AM9624 |
| 1× PBS | Thermo Fisher Scientific | Cat. # BP24384 |
| Nuclease-free water | Thermo Fisher Scientific | Cat. # AM9932 |
| Low melt agarose | IBI Scientific | Cat. # IB70056 |
| Ethyl alcohol (EtOH) | MilliporeSigma | Cat. # E7023 |
| MOPS | MilliporeSigma | Cat. # M1254 |
| Melphalan | Cayman Chemical | Cat. # 16665 |
| Acryloyl-X, SE (AcX) | Thermo Fisher Scientific | Cat. # A20770 |
| DMSO (anhydrous) | Thermo Fisher Scientific | Cat. # D12345 |
| Poly-L-Lysine | TED PELLA, INC | Cat. # 18026 |
| Photo Flo 200 | Electron Microscopy Sciences | Cat. # 74257 |
| 40% Acrylamide | MilliporeSigma | Cat. # A4058 |
| 2% *N,N'*-Methylenebisacrylamide | MilliporeSigma | Cat. # M1533 |
| Acrylic acid (anhydrous) | MilliporeSigma | Cat. # 147230 |
| 10 N Sodium hydroxide (NaOH) | Thermo Fisher Scientific | Cat. # SS267 |
| *N,N,N',N'*-Tetramethyl ethylenediamine (TEMED) | MilliporeSigma | Cat. # T7024 |
| Ammonium persulfate (APS) | MilliporeSigma | Cat. # A3678 |
| 4-Hydroxy-TEMPO (4-HT) | MilliporeSigma | Cat. # 176141 |
| Proteinase K (ProK) | New England Biolabs | Cat. # P8107S |
| 1 M Tris (pH8) | Thermo Fisher Scientific | Cat. # AM9855G |
| 0.5 M EDTA | Thermo Fisher Scientific | Cat. # AAJ15694AE |
| 5 M NaCl | Thermo Fisher Scientific | Cat. # AM9760G |
| 10% SDS | VWR | Cat. # 97062-964 |
| Triton X-100 | Thermo Fisher Scientific | Cat. # AAA16046-AE |
| RNase-Free DNase I | Qiagen | Cat. # 79254 |
| 1 M MgCl<sub>2</sub> | VWR | Cat. # 97062-848 |
| CaCl<sub>2</sub> | VWR | Cat. # 97062-590 |
| Formamide | Thermo Fisher Scientific | Cat. # BP227-100 |
| 20× SSC | Thermo Fisher Scientific | Cat. # AM9763 |
| 50% Dextran sulfate | MilliporeSigma | Cat. # S4030 |
| Tween 20 | Thermo Fisher Scientific | Cat. # AAJ20605AP |
| Probe Hybridization Buffer | Molecular Instrument | https://store.molecularinstruments.com/new-kit/v3/rnafish |
| Probe Wash Buffer | Molecular Instrument | https://store.molecularinstruments.com/new-kit/v3/rnafish |
| Amplifier Buffer | Molecular Instrument | https://store.molecularinstruments.com/new-kit/v3/rnafish |
| DAPI | MilliporeSigma | Cat. # D9542 |
| RNase AWAY™ Decontamination Reagent | Thermo Fisher Scientific | Cat. # 10328011 |
| **Experimental models: Organisms/strains** |  |  |
| Mouse: C57BL/6J | Jackson Laboratory | JAX stock #000664 |
| Mouse: Thy1-GCaMP6f (GP5.17Dkim/J) | Jackson Laboratory | JAX stock #025393 |
| **Oligonucleotides** |  |  |
| HCR probes | Molecular Instrument | N/A |
| HCR Amplifier B1, B2, B3, B4, B5 | Molecular Instrument | N/A |
| HCR Amplifier with JF669 | Wang et al., 2021 | N/A |
| **Software and algorithms** |  |  |
| EASI-FISH pipeline | Wang *et al.*, 2021 | https://doi.org/10.5281/zenodo.5652239<br>https://janeliascicomp.github.io/multifish/ |
| Fiji | National Institute of Health | https://imagej.net/software/fiji/ |
| Python |  | Python.org; RRID: SCR 008394 |
| BigWarp plugin for Fiji | Bogovic, J.A. *et al.*, 2016 | https://imagej.net/plugins/bigwarp |
| BigStream |  | https://github.com/JaneliaSciComp/bigstream |
| Suite2p | Pachitariu *et al.*, 2017 | https://github.com/MouseLand/suite2p |
| **Other** |  |  |
| Zeiss Lightsheet Z.7 microscope | Zeiss | https://www.zeiss.com/microscopy/en/products/light-microscopes/light-sheet-microscopes/lightsheet-7.html |
| Press-to-Seal™ Silicone Isolator with Adhesive, 9 mm diameter, 0.5 mm deep | Thermo Fisher Scientific | Cat. # P24743 |
| 8mm glass coverslip | Harvard Apparatus | Cat. # BS4 64-0701 |
| Zeiss Lightsheet Z.7 imaging holder |  | https://hhmi.flintbox.com/technologies/65795945-938b-4115-b2ed-8ccfb9b2e511 |

## EXPERIMENTAL MODEL AND STUDY PARTICIPANT DETAILS

### Mice

Adult male and female transgenic mice (Thy1-GCaMP6f59 (GP5.17Dkim/J), 8-12 weeks of age at the beginning of the experiments were used. All methods for animal care and use were conducted according to National Institutes of Health guidelines for animal research and approved by the Institutional Animal Care and Use Committee (IACUC) at Cornell University and at Janelia Research Campus. Mice were housed in a 12 h light/12 h dark cycle and had *ad libitum* access to water and chow diet.

## METHOD DETAILS

### Surgery

All animals were anesthetized with 1.5% isoflurane (induction phase rate at 3%), with the top fur of the head shaved, and then placed in a stereotaxic frame with a heating pad underneath. Eye ointment was applied and lidocaine (7 mg/kg, s.q.) and bupivacaine (5 mg/kg, s.q.) were injected under the shaved area. The shaved area was cleaned with cotton swabs with 70% ethanol and betadine. Then, an incision was made, and the skull was exposed to level the bregma and lambda as well as medio-lateral axis. A craniotomy on the right hemisphere was performed, centered at 3.1 mm anteroposterior and 3.0 mm mediolateral from the bregma using a 3-mm diameter trephine drill bit. Once the skull was removed, dura was carefully removed with the tip of a fine needle. From this point until the cannula was placed, ice-cold sterile cortex buffer ^75^ was applied to prevent the tissue from drying and to flush out blood. The cortex overlying the dorsal hippocampus was gently aspirated using a 25-gauge blunt-tip needle connected to a vacuum system until the fibers of the corpus callosum became visible. A 3-mm glass coverslip, previously attached to a stainless-steel cannula with optical glue, was placed over the dorsal hippocampal region. The cannula and a custom titanium headbar were secured to the skull with dental cement. After surgery, all animals received ketoprofen (5 mg/kg, *s.q*.) daily for three days and were allowed to recover for a minimum of one week before starting water restriction (1.0-1.5 mL daily) in a reversed dark-light cycle room (12 h light/12 h dark cycle).

### Behavior and Calcium imaging

Behavioral procedures were as previously described ^47^. Briefly, mice were placed on water restriction (1.0-1.5 mL water) for at least 2 weeks before behavior training. For behavior, mice were acclimated to run on a spherical treadmill with head-fixation and then trained to learn a head-fixed virtual-reality task (the two-alternative cue-delay-choice task) in which the indicator cues predicted reward availability, with one cue associated with water reward at a near location and the other cue associated with water reward at a far location. Water rewards were obtained by licking at the appropriate reward zone. For the first 3 days, guided training was performed, where water rewards were delivered at the correct location even without licking at the reward zone. At the end of the guided training, consistent anticipatory lickings were observed. For subsequent training, water reward was delivered only when mice licked in the correct reward zone. No penalty was imposed for licking at the incorrect locations. Each trial type was randomly selected with 50% probability, with a maximum of three consecutive repeats. Throughout training, neural activity was recorded using the two-photon random access (2P-RAM) mesoscope ^46^ and data was acquired at a scanning frequency of 10 Hz using ScanImage software. The imaging data was analyzed using the Suite2p toolbox (https://www.suite2p.org/), including motion correction, segmentation, neuropil correction and spike deconvolution as described elsewhere ^50,51^.

### Neural data analysis

#### Calcium event detection

Calcium transients were identified as previously described ^47^. For each cell, the ΔF/F₀ within the speed-filtered running window were calculated based on neuropil-corrected traces. ΔF/F_0_ was defined as the difference between fluorescent and baseline activity divided by F_0_. F_0_ for each cell was calculated by first applying Gaussian filter (5 s) followed by calculating the rolling max of the rolling min (‘maximin’ filter). Calcium transient event was defined as events that started when fluorescence deviated 5σ from baseline and ended when it returned to within 1σ of baseline. σ was calculated by binning the fluorescent trace in short periods of 5 s and considering only frames with fluorescence in the lower 25th percentile. Event amplitude was defined as the peak fluorescence during the event duration. Event rate was computed as the number of calcium transients within the running window divided by the duration of that window in seconds (events/s).

#### Spatial information and stability

Spatial information (SI) was computed for each cell using the Skaggs formula ^76,77^, also used in ^78^. SI = Σᵢ pᵢ (rᵢ/R)log₂(rᵢ/R), where rᵢ is the trial-averaged smoothed ΔF/F₀ in spatial bin i (bin size: 5 cm), R = Σᵢ pᵢ rᵢ is the occupancy-weighted mean ΔF/F₀, and pᵢ is the measured fractional occupancy of bin i (time spent in bin i / total time, computed from speed-filtered position frames). Between-session stability was computed as the Pearson correlation coefficient between the cell’s trial-averaged rate map in session N and its trial-averaged rate map in session N − 1, computed separately for each reward condition (cells were compared under matched reward conditions on consecutive days). Rate maps were derived from the trial-averaged ΔF/F₀ across spatial bins (5 cm bins, 230 cm linear track) ^79^.

Session 0 was not included due to lack of preceding session. For within-session stability, the Pearson correlation was performed on the rate maps from odd- and even-trials for each session and each reward type. Between-session stability measures persistence of the spatial code across days whereas within-session stability measures reliability of the spatial map between trials.

#### Place field detection

Place fields are detected as previously described ^47^ during active trials. First, putative place fields were identified by spatially binning the resulting ΔF/F_0_ activity (bin size of 5 cm) as continuous regions where all ΔF/F_0_ values exceeded 25% of the difference between the peak of the trial and the baseline 25th percentile ΔF/F_0_ values. Additional criteria were imposed: the field width should be between 15 and 120 cm in virtual reality, the average ΔF/F_0_ inside the field should be at least four times greater than outside; and significant calcium transients should occur at least 20% of the time when the mouse was active within the field. To verify that these putative place fields were not caused by spurious activity, we calculated a shuffle-based null distribution for each cell. Calcium activity was shuffled in 10 s blocks with respect to the animal’s position, and the same place-field detection procedure was applied to the shuffled data. This process was repeated 1,000 times and a cell was considered to have a significant place field if putative fields were detected in fewer than 5% of shuffles.

All metrics were averaged across sessions and reward types for cell cluster comparisons and molecular gradient analyses.

### Acquisition of high-resolution *in vivo* z-stack images

At the end of the last *in vivo* imaging session, a high-resolution z-stack covering the two-photon imaged region was taken before the animal was transcardially perfused. To do this, the parameters were set as follows: lateral resolution = 1 µm, z step size = 4 µm, averaging every 10 frames, z-depth 50-100 µm spanning dorsally and ventrally from the imaging plane. The z-depth from the dorsal tissue surface to the imaging plane was documented to facilitate subsequent identification of the *in vivo* plane in *ex vivo* tissue volume.

### Tissue fixation and preparation

All procedures were performed in RNase-free environment. Animals were anesthetized with isoflurane and perfused with RNase-free DPBS followed by freshly prepared ice-cold 4% paraformaldehyde (PFA). After carefully removing the cranial window, the brain tissue was dissected and fixed in 4% PFA overnight at 4°C before sectioning on a vibratome (**Figure 1B**). Brains were sectioned horizontally at 300 μm thickness and stored in 70% ethanol at 4°C.

### EASI-FISH with GCaMP imaging protocol summary

EASI-FISH for multi-round detection of RNA was carried out as previously described ^32^ with an additional step to image GCaMP for the *in vivo* to *ex vivo* registration. In brief, RNA and proteins in the tissue were covalently anchored to the hydrogel using MelphaX and AcX, respectively. Tissue was then embedded into the hydrogel, followed by proteinase K digestion (500mM NaCl). The digested sample was then treated with DNase I to remove genomic DNA, allowing DAPI to produce cytosolic staining (*i.e*., cytoDAPI), which was used for cell segmentation and multi-round EASI-FISH registration. At this point, the sample was imaged with the Lightsheet microscope to obtain Round 0 (GCaMP and cytoDAPI) z-stack, which served as bridging data between *in vivo* and *ex vivo* (**Figures. 1C, left and 2A**). After imaging, the sample was incubated in disruption buffer (50 mM Tris, pH 8.0, 200 mM NaCl, and 5% SDS) for 1 h at 60°C, followed by overnight Proteinase K treatment (50mM NaCl) at 37°C, to eliminate GCaMP fluorescence and free the green channel for subsequent EASI-FISH (**Fig. 1C, middle and right**). Marker genes used for EASI-FISH in the dorsal CA1 include *Cbln4*, *Col11a1*, *Ly6g6e*, *Lefty1*, *Syt2*, *Nts*, *Necab1*, *Fn1*, *Spink8*, *Klk8*, *S100b*, *Fndc5*, *Lpl*, *Dlk1*, *Tpbg*, *Ndst4*, *Tshz2*, *Lrp1b*, *Fxyd6*, and *Snap25*. Marker gene expressions were detected using HCR probes and hairpins ^80^. Samples were imaged on a Zeiss Lightsheet Z.7 microscope with a 20× water-immersion objective (20×/1.0 W Plan-Apochromat Corr DIC M27 75 mm, RI = 1.33) and analyzed using the EASI-FISH computational pipeline as previously described ^32^. DNase I digestion was used for round-to-round stripping followed by cytoDAPI staining to generate the cytosolic staining pattern for registration and segmentation.

### *Ex vivo* to *in vivo* image registration

The *in vivo* z-stack (voxel size of 1 × 1 × 4 µm³ in x, y, and z) served as the fixed image, and the *ex vivo* EASI-FISH image stack (s3 scale, voxel size of 0.93 × 0.93 × 0.84 µm³ in x, y, and z) served as the moving image for registration. The registration consisted of two stages, global affine alignment followed by distributed block-wise deformable registration. First, corresponding anatomical landmarks pairs (n=30-50) were manually identified in both the fixed and moving images using BigWarp ^81^ in Fiji/ImageJ. Landmark coordinates were exported in physical units. A 3D affine transformation was estimated from matched landmark pairs using RANSAC-based robust fitting implemented in OpenCV (cv2.estimateAffine3D). To correct local nonlinear tissue distortions, an open-source Python toolkit for large-scale volumetric image registration called BigStream ^38^ was used. Images were first intensity-normalized and then partitioned into overlapping blocks (block size: 256 × 256 × 128 voxels; overlap fraction: 0.5). Each block was initialized with the global affine transform, then the B-spline deformable registration was optimized based on ANTs neighborhood correlation ^82^ implemented in SimpleITK. The *in vivo* z-stack (voxel size of 1 × 1 × 4 µm³ in x, y, and z) served as the fixed image, and the *ex vivo* EASI-FISH image stack (s3 scale, voxel size of 0.93 × 0.93 × 0.84 µm³ in x, y, and z) served as the moving image for registration. The registration consisted of two stages, global affine alignment followed by distributed block-wise deformable registration. First, corresponding anatomical landmark pairs (n=30-50) were manually identified in both the fixed and moving images using BigWarp ^81^ in Fiji/ImageJ. Landmark coordinates were exported in physical units. A 3D affine transformation was estimated from matched landmark pairs using RANSAC-based robust fitting implemented in OpenCV (cv2.estimateAffine3D). To correct local nonlinear tissue distortions, an open-source Python toolkit for large-scale volumetric image registration called BigStream ^38^ was used. Images were first intensity-normalized and then partitioned into overlapping blocks (block size: 256 × 256 × 128 voxels; fraction of overlap: 0.5). Each block was initialized with the global affine transform, then the B-spline deformable registration was optimized based on ANTs neighborhood correlation ^82^ implemented in SimpleITK. Then the block-wise deformation fields were stitched with linear blending into a single displacement vector field.

Registration performance was assessed at different stages using four complementary metrics. Target registration error (TRE) was calculated as the Euclidean distance between corresponding hold-out manually annotated landmark pairs, with 30-80 landmarks used per image pair. Normalized cross-correlation (NCC) was computed as the global Pearson correlation coefficient between normalized fixed and moving images. Local cross-correlation was calculated as the spatial mean of the cross-correlation between fixed and moving images within an 11×11×11-voxel sliding window (radius 5). Mutual information (MI) was computed from the joint and marginal intensity histograms of the fixed and moving images.

### Cell mask matching

The mean fluorescence field-of-view (FOV) image and cell masks produced by Suite2P were registered to the *in vivo* z-stack coordinate system using a two-step pipeline. First, a global 2D affine transformation was estimated using SIFT feature matching ^83^ between the Suite2P FOV image and the target z-plane, with RANSAC used to remove outlier feature matches. Second, deformable registration implemented in BigStream was applied to correct for local deformations. The Suite2P cell masks and ex vivo cell masks were then warped into the *in vivo* z-stack coordinate system using nearest-neighbor interpolation to preserve integer cell labels. For each *in vivo* Suite2P cell, candidate *ex vivo* matches were identified within the target z-plane ± 3 planes based on spatial overlap with warped *ex vivo* cell masks. Each candidate pair was characterized by intersection-over-union (IoU), centered IoU (cIOU, IoU computed after centering both masks on their centroids), and Euclidean centroid distance in physical units. To establish a matching criterion, candidate pairs were stratified and sampled across the cIoU-distance feature space and manually labeled as true matches, false matches, or ambiguous. A logistic regression classifier was fit using standardized cIoU and centroid-distance features from the labeled candidate pairs, and the operating threshold p* was selected by maximizing Youden’s index (J = sensitivity + specificity − 1). The trained classifier (Sensitivity: 0.92, Specificity: 0.91) was then applied to all candidate pairs to estimate match probability, and final matches were assigned greedily under a one-to-one constraint: each *in vivo* cell matched to at most one *ex vivo* cell across all z-planes, ranking candidates by predicted probability. Candidate pairs with predicted probability ≥ p* were retained for downstream analysis. Pipeline precision was assessed by manually inspecting 50 randomly selected pairs from the final accepted one-to-one matched pool. Precision and 95% confidence interval were reported.

### EASI-FISH data analysis

Image processing, including stitching, registration, segmentation, and spot extraction, as well as cell clustering and UMAP embedding were performed as previously described ^32^. Principal component analysis (PCA) was applied to clusters C1 and C2 to extract the molecular gradient. Gene count was normalized, log-transformed, and then z-scored for PCA analysis. The first principal component (PC1) was used as the molecular gradient in all subsequent analyses.

## QUANTIFICATION AND STATISTICAL ANALYSIS

Non-parametric Mann-Whitney U tests were used to compare cell clusters and p values were reported. Associations between the molecular gradient (PC1) and functional properties, including calcium transient measure and spatial coding properties, were assessed using Spearman’s rank correlation. Spearman’s correlation coefficient, ρ was reported as a measure of effect size, and the corresponding p value was reported to assess statistical significance. To examine whether the relationship between PC1 and functional metrics changed across learning, sessions were classified into four training stages based on behavioral performance: pre-training (guided sessions), early (success rate <50% for both trial types), middle (success rate >50% for trial type 1 and <50% for trial type 2), and late (success rate >80% for both trial types). Kruskal– Wallis tests were used to compare changes across training stages. Additional quantitative analyses in the study are described in the relevant method sections above.

